# Dementia risk genes engage gene networks poised to tune the immune response towards chronic inflammatory states

**DOI:** 10.1101/597542

**Authors:** Jessica Rexach, Vivek Swarup, Timothy Chang, Daniel Geschwind

## Abstract

An emerging challenge in neurodegenerative dementia is understanding how immune-associated genes and pathways contribute to disease. To achieve a refined view of neuroinflammatory signaling across neurodegeneration, we took an integrative functional genomics approach to consider neurodegeneration from the perspective of microglia and their interactions with other cells. Using large-scale gene expression and perturbation data, regulatory motif analysis, and gene knockout studies, we identify and characterize a microglial-centric network involving distinct gene co-expression modules associated with progressive stages of neurodegeneration. These modules, which are conserved from mouse to human, differentially incorporate specific immune sensors of cellular damage and pathways that are predicted to eventually tune the immune response toward chronic inflammation and immune suppression. Notably, common genetic risk for Alzheimer’s disease (AD), Frontotemporal dementia (FTD) and Progressive Supranuclear Palsy (PSP) resides in specific modules that distinguish between the disorders, but also show convergence on pathways related to anti-viral defense mechanisms. These results suggest a model wherein combinatorial microglial-immune signaling integrate specific immune activators and disease genes that lead to the establishment of chronic states of simultaneous inflammation and immunosuppression involving type 1 interferon in these dementias.

## Introduction

Leveraging genetic discoveries to identify therapeutic targets requires understanding how disease genes map onto cell-type specific molecular pathways. In this regard, a remarkable body of growing genetic evidence supports a link between Alzheimer’s and associated dementias and neuroimmune functions involving glial cells^1,2^ Causal genetic variation in neurodegenerative dementias affects genes with neural-immune functions affecting two major CNS resident neural immune cells, astrocytes and microglia, including, *TREM2, GRN, HLA-DRA, CR1, C9ORF72, APOE, BIN1, CXCR4, CLU*, and TBK1^1–6^. Multiple functional studies in animal models of neurodegeneration support the contribution of microglial and neural-immune genes to disease-associated phenotypes including age-associated cognitive decline, pathological protein deposition and dyshomeostasis, and neurodegeneration^7–9^. The discovery that immune-related genes contribute to AD and associated dementias has generated great enthusiasm for the possibility of immune-based therapies^10^. From this perspective, defining the detailed molecular relationship between disease-associated neuroimmune pathways and causal dementia genes has the potential to inform disease mechanism and inspire novel therapeutic approaches.

Microglia and CNS-resident macrophages are the principle immune cells of the brain with critical roles in detecting immunogens and coordinating the immune response^11^. During nervous system injury, microglia can be directly activated by myelin, lipids, or nucleotides released from injured cells to activate pro-inflammatory signaling, such as through the NLRP3 inflammasome complex^12,13^. Experimental evidence in multiple models of AD pathology suggests that “disease-associated microglia” express dementia risk genes, including *APOE* and *TREM2*, and contribute to synaptic injury, neurotoxic astrocyte transitions, and neuronal dysfunction^14–17^. Furthermore, single-cell genomic studies have begun to delineate heterogeneity among disease-associated microglial states and their trajectories, highlighting the need to better understand their specific roles in neurodegeneration^18,19^. This includes defining how different microglial states relate to disease-associated immune activators and causal genes in human neurodegenerative diseases.

We recently used a systems biology approach to integrate gene expression data from human post mortem brain and multiple mouse models harboring human dementia causing mutations, to identify a robust neurodegeneration-associated inflammatory module (NAI) and a closely correlated neurodegeneration-associated synaptic module (NAS)^20^. The NAI module is strongly enriched for markers of both astrocytes and microglia, both of which are known to be significantly up-regulated in multiple neurodegenerative syndromes^9,21,22^. As a result of this global up-regulation within tissue, the cell-type specific expression patterns of glial genes in the NIA module were obscured. In silico approaches for deconvoluting cell-specific signatures are challenged by the complex dynamics among glial genes in disease^16,18,19,23,24^. So, we reasoned that the optimal resource for resolving glial pathways involved in neurodegeneration would be gene expression data from actual glial cell types isolated from disease and control samples. Furthermore, given that neurodegeneration involves interactions between neurons and among glia^25^; we reasoned that integrating data from sorted cells and intact tissue would reveal disease-relevant and cell-specific signaling networks.

Here we present an integrative analysis of microglial-specific transcriptomic changes that are latent components of neurodegeneration pathways at the tissue-level. Our findings parse disease genes into distinct microglial co-expression sub-networks (modules) related to progressive stages of neuropathology in mice that are conserved in humans. Using large-scale gene perturbation data, regulatory motif analysis, and knockout studies, we identify strong evidence for regulatory interplay that functionally connects different modules into a microglial-centered interactome. By incorporating genetic association data, we find that the genetic risk factors contributing to Alzheimer’s disease (AD), Frontotemporal dementia (FTD) and Progressive Supranuclear Palsy (PSP) involved shared and distinct microglia-associated neuroimmune modules. However, as disease progresses, the associated shared transcriptional and PPI networks that are up-regulated involve chronic viral response pathways to double stranded RNA, likely driven by Type-1 interferon, supporting a model whereby early immune activation gives way to chronic immunosuppression in these disorders.

## Results

We performed consensus weighted gene co-expression analysis (WGCNA^26^; **Methods**) to combine gene expression data from sorted, purified microglia from a mutant tau mouse model (rTg4510; AMP-AD Knowledge Portal doi:10.7303/syn2580853) and whole brain tissue collected from multiple independent transgenic mouse models of neurodegenerative tauopathy (**Methods, Figure 1A**) to identify conserved modules that exist in both purified microglia and tissue. We then used the microglia-specific gene expression data to identify up and down-regulated pathways (**Figure 1A, Schema**). Using this approach, we identified 13 distinct co-expression modules varying in disease association, trajectory and time course (**Fig 1B, 1C, 1D, 2A**).

**Figure 1:**
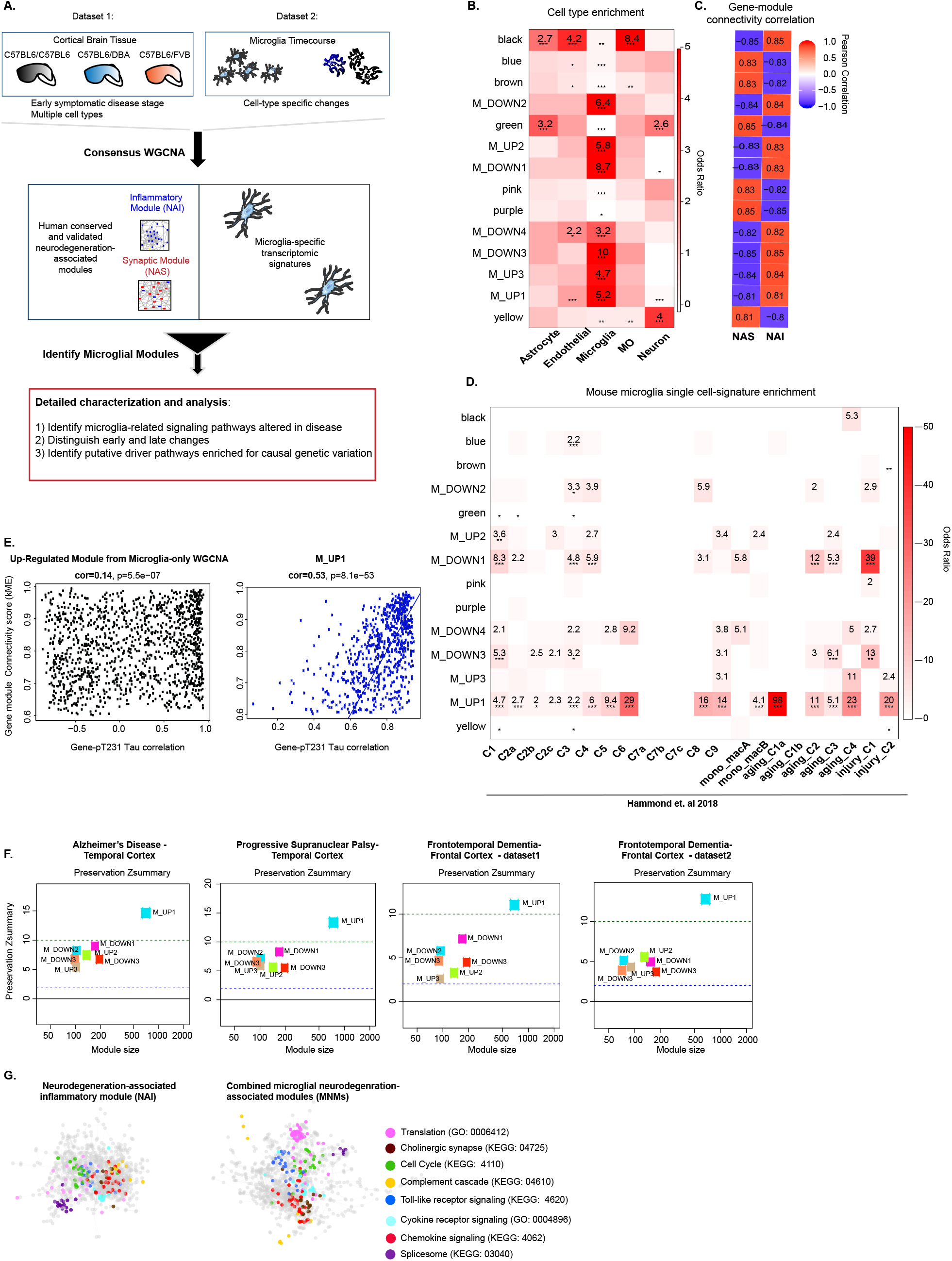
Purified microglia-brain tissue consensus gene co-expression network analysis. **A**, Experimental schema, showing approach for microglia-tissue consensus WGNCA and module selection to achieve multiple microglial disease modules associated with tissue-level neurodegenerative disease inflammatory modules. **B**, Cell type enrichment of modules using mRNA markers for corresponding cell types from mouse brain (Fisher’s two-tailed exact test, ***FDR<0.005, MO = myelinating oligodendrocyte^27^). **C**, Pearson’s product-moment correlation (n = 12413 genes) of module eigengene connectivity in consensus module compared to tissue-level neurodegeneration module (NAS or NAI)^20^. **D**, Module enrichment for mouse microglia single cell cluster signatures (as defined in^28^; Fisher’s two-tailed exact test, *FDR<0.05, **FDR<0.001, ***FDR<0.005). **E**, Scatterplot showing Pearson’s correlation of gene-module connectivity (kME) and sample-by-sample correlation of gene expression and pT231 Tau levels (n=36) in TPR50 mouse brain (frontal cortex, 6 months of age, n=18 per group of WT or P301L MAPT; P-values obtained from two-sided test for Pearson correlation are shown)^20^. **F**, Module preservation in human AD and control temporal cortex (control n=308, AD n =157)^33^, human PSP and control temporal cortex (control n=73, PSP n =83)^33^, and human FTD and control frontal cortex from two independent datasets (dataset 1^95^ control n=14, FTD n=16; dataset 2^20^: control n=8, FTD n=10). The bottom line is at the lower cut off for preservation (Zsummary = 2) and the upper line in at the cut off for high preservation (Zsummary = 10) as defined in^84^. **G**, Protein-protein interaction (PPI) network plot of among all genes from tissue-level NAI (left) and combined microglia-enriched consensus modules (MNMs; right), with nodes colored by GO and KEGG categories, as shown.

### Consensus microglial modules combine cell-type specificity and tissue-level neuronal-glial relationships

We first focused on the 7 modules significantly enriched for genes expressed in microglia compared to other cell types^27^ (**Fig. 1B, 1D; Supplementary Fig. 1A**). As independent validation of cell-type trends, we assessed module enrichment for single cell microglial signatures, previously identified from high resolution single cell sequencing studies in mouse^28^ and human^29^ brain, and observed significantly greater marker enrichment among these 7 candidate microglia-gene enriched modules compared to the remaining modules (**Fig. 1D, Supplementary Fig. 1A**). At the same time, these modules were distinct in that different modules overlapped with different sets of microglial signatures identified in previous single cell analyses^28,29^ suggesting that they represented distinct microglia pools (**Fig. 1D, 2E, Supplementary Fig. 1A, 1G**). For example, M_UP1 enriched for profiles of microglia proliferative states (clusters 2a, 2b, 2c)^28^ and age-associated states identified previously in single cell transcriptome analyses (C8, aging_C1, aging_C2, aging_C3)^28^ (**Fig. 1D**).

**Figure 2:**
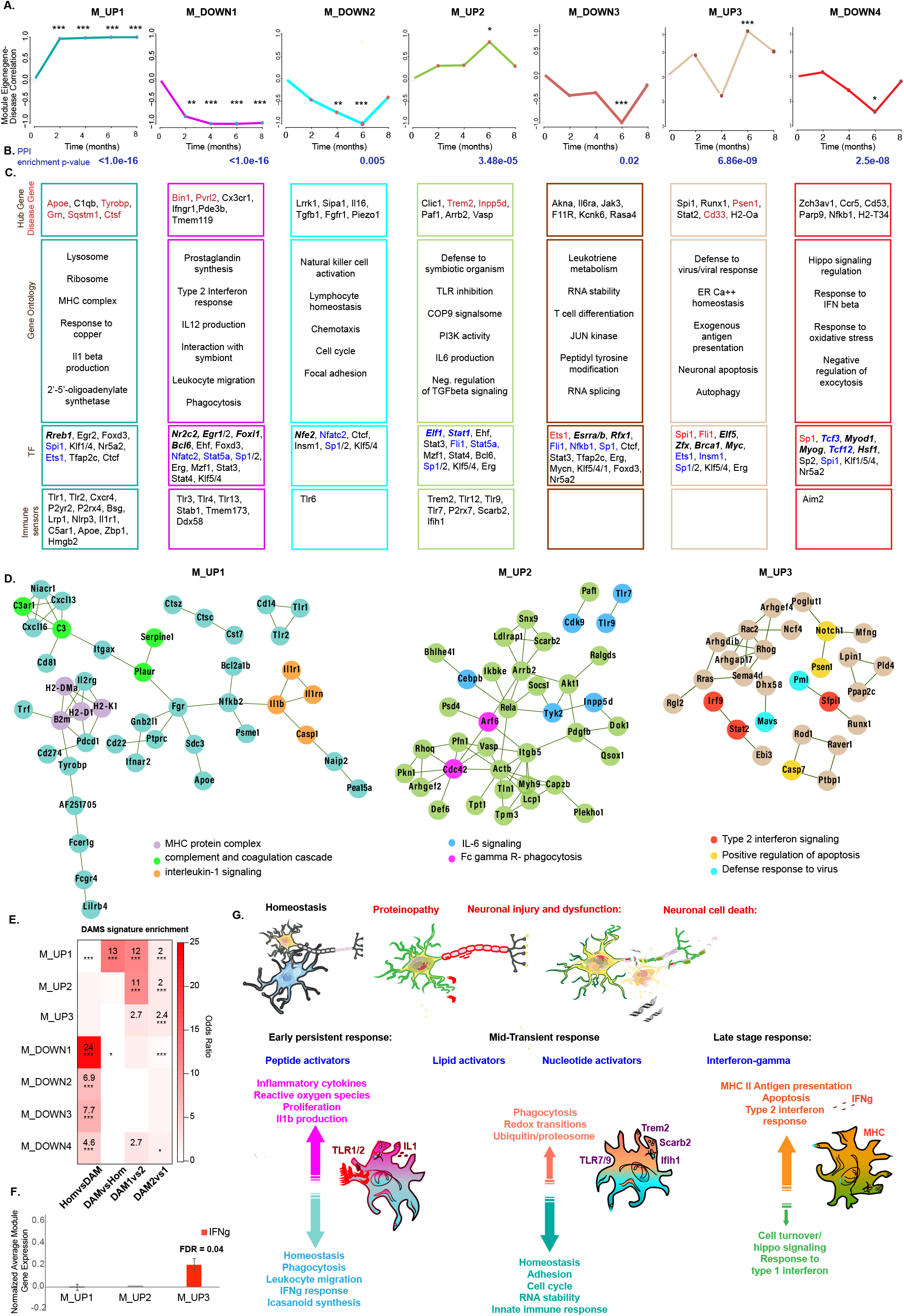
Microglia-tissue consensus module microglia disease time-course and pathway annotation. **A**, Signed Pearson’s correlation of the module eigengene (ME) calculated in the rTg4510 microglia gene expression dataset at each age (unpaired two-tailed T-test; n=7 modules, n=4 mice per genotype (P301L MAPT or WT) per timepoint; *p-value<0.05, **p-value<0.01, ***p-value<0.005). Graphed with theoretical zero plotted at time zero. **B**, Module PPI network enrichment p-value (p-value calculated as described in^96^). **C**. Select module genes (with disease genes in red), enriched gene ontology terms (Z-score >2), transcription factors (TF) with binding site enrichment (labels are bold and italic if the TF is unique to one module, blue if the TF is a hub gene in any module, and red if the TF is a hub gene in the same module; p-value <0.05 compared to whole genome CpG islands), and module genes that are receptors of pathogen or damage associated molecular patterns (“immune sensors”). **D**. Protein-protein interactions among top 150 module genes (ranked by kME) with enriched pathway genes labeled (GO-Elite^83^ permuted Z score >2). **E**, Module enrichment heatmap for top 100 genes differentially expressed between progressive microglia single cell states, as indicated, Hom = homeostatic, DAM1 = type 1 disease-associated microglia (DAM), DAM2 = type 2 DAMs, as defined in^18^; n=7 modules with 4 comparisons per module, *FDR <0.05, **FDR<0.005, ***FDR<0.001. **F**, Differential module expression in purified microglia following treatment with IFN-gamma (n=8) compared to untreated controls (n=4) (two-tailed unpaired T-test, human fetal microglia cells, IFNg at 200u/mL for 6h or 24h, GSE143 2^41^). **G**, Model showing microglia transitions across progressive disease stages based on annotation of microglia-tissue consensus modules (MNMs).

We next tested whether our consensus modules recapitulate biological relationships present in tissue-level neurodegeneration modules identified in prior studies (NAI and NAS)^20^. As expected, the seven microglia-enriched modules were highly positively correlated with the NAI inflammatory module and negatively correlated with the NAS synaptic module; both of which have been previously demonstrated to be conserved across humans with tauopathies and mice harboring mutations causing dominant forms of FTD in humans^20^ (**Fig. 1C**).

Next, we assessed each module’s relationship to pathological neuronal Tau hyperphosphorylation, a measure of neuropathology associated with disease progression^30^. We found a strong positive correlation between pathological Tau phosphorylation levels and microglia-enriched consensus module gene connectivity (**Fig. 1E, Supplementary 1B**). In contrast, when we analyzed WGCNA modules generated using only sorted microglial cell gene expression data from the rTg4510 model, rather than consensus modules based on network edges shared between whole tissue and the sorted cell data, the correlation with pTau was substantially reduced (**Fig. 1E**). This demonstrates the utility of using both cell specific and whole tissue data to advance our understanding of cell specific contributions to disease pathology.

Finally, we observed that the NAI and combined microglial consensus modules are conserved at the level of protein-protein interactions (PPI), which themselves coalesce into distinct molecular pathways (**Fig. 1G, Supplementary Fig. 1C**). Thus, these seven co-expression modules represent a substantial refinement of a previously identified neural immune module observed in both model systems and post mortem human brain^18–20,31,32^. We label the seven new modules “microglia-associated neurodegeneration-associated modules” (MNMs, 1-7), and further characterized them as a means to explore associated microglial functional pathways and regulators related to neurodegeneration.

### MNMs are conserved in human disease brain and mouse models

To assess the robustness of the microglia-enriched modules and further validate their relevance to human disease, we tested their preservation using multiple independent mouse and human disease datasets (see **Methods**). Consistent with the observation of PPI conservation for all modules, all seven MNMs are preserved in post mortem human brain tissue from AD^33^, FTD^20,34^ and PSP^33^ patients (**Fig. 1F**). Additionally, all MNMs are preserved in three different transgenic mouse models expressing human MAPT mutations^20,35^ (**Supplementary Fig. 1D**) and in microglial-specific datasets from mouse models expressing PSEN^36^ and APP^37^ mutations, except M_UP3, which is only weakly preserved in one of two datasets (**Supplementary Fig. 1E**). However, we do note that the differential expression patterns of three modules (M_UP2, M_UP3, M_DOWN3) differ between microglia isolated from P301L MAPT (rTg4510) and PSEN/APP mutant mouse models (**Supplementary Fig. 1F**). Together, the preservation of these modules across multiple independent disease datasets including human disease brain from Alzheimer’s and associated dementias and mouse models of Alzheimer’s or FTD, as well as PPI, indicates that they represent robust biological processes. However, some modules display variability in their differential expression in different disease models, suggesting they may be conditional on disease-stage or disease-specific pathology.

### Microglial molecular transitions along progressive epochs of neuronal pathology

In contrast to the composite whole tissue NAI module expression vector which shows a singular increasing trajectory over time (shown in^20^), we were able to deconvolute the MNM modules into highly distinct temporal trajectories with respect to progressive disease stages modeled in the rTg4510 mouse between 2 and 8 months of age^35,38–40^. We identified three temporal patterns of module-disease association: (1) changing at the earliest disease stage, prior to neuronal loss, and persistent through later stages (M_UP1, M_DOWN1, M_DOWN2), (2) changing during early periods of neuronal loss, and more transient (M_UP2, M_DOWN3), and (3) most significant changes during late stages of continued neuronal loss and cumulative pathology (M_UP3) (**Fig. 2A**). Therefore, combined tissue-microglial cell consensus WGCNA resulted in microglial neurodegeneration-associated modules with distinct temporal transitions across disease progression, which were present in latent forms, but not detected in the analysis of whole tissue alone.

To validate our MNM-disease stage trends using complementary published datasets, we compared them to time-course, single-cell microglial differential gene expression data collected from two different mouse models with Alzheimer’s related pathology (5xFAD^18^, CK-p25^19^), including one model with frank neurodegeneration (CK-p25^19^). As expected, the early up-regulated MNM, M_UP1, is enriched for genes that are increased in early microglial disease states relative to homeostatic microglia (**Fig. 2E, Supplementary Fig. 1G**), and the later up-regulated MNMs are enriched for genes up-regulated in later relative to early microglial disease states (**Fig. 2E, Supplementary Fig. 1G**). The four down-regulated MNM are all enriched for microglial genes down-regulated in early microglia disease states (**Fig. 2E, Supplementary Fig. 1G**).

We next leveraged published data on the type 2 interferon response to ask whether MNMs reproduce the late interferon-gamma signature reported to distinguish microglia during periods of neuronal cell death^19^. We were able to show that the last up-regulated MNM in disease, M_UP3, is also the only module induced by interferon-gamma treatment of cultured microglia^41^ (**Fig. 2F**), consistent with the published trend^19^. Altogether, these findings support that MNMs recapitulate stage-associated, microglia-specific biological trends identified from recent single-cell studies using mouse models of Alzheimer’s pathology^18,19^. Moreover, MNMs further refine prior findings, separating disease-associated microglia changes across multiple distinct modules. Therefore, we next explored these MNMs in detail to delineate stage-associated transitions in microglia signaling including changes prior to and subsequent to cell death, and their relationship to dementia disease genes.

### Pathway analysis to expand biological insights into microglial transitions across disease

Annotation of modules for enriched biological regulators and pathways aligned specific disease genes, signaling and functional pathways, immune receptors, transcription factors and microglia-enriched gene co-expression modules with progressive stages of neuronal dysfunction and degeneration that are summarized in Figure 2 (**Fig. 2A, 2B, 2C, 2D, 2F, 2G, Supplementary Fig. 2A, 2B**). These annotations indicate that each of these modules represents different aspects of the microglia function that vary across disease stage, with different microglial modules poised to sense and respond to specific damage-associated immune activators^12,42^ that change over time (**Fig. 2A, 2C, 2G**).

For example, the earliest up-regulated module, M_UP_1, incudes sensors of peptide and lipopeptide immune activators (TLR1, TLR2), whereas the subsequently up-regulated module, M_UP2, includes sensors of lipid immune activators (TREM2, SCARB2) together with receptors for viral nucleotides (TLR7, TLR9, Ifih1) that can also be activated by damaged or dysregulated endogenous DNA^12,42–46^. Therefore, our microglial time-course analysis shows a prominence of DNA and RNA detecting immune receptors within the second phase of up-regulated MNMs, suggesting that nucleic acids activate inflammatory pathways as neuronal injury and disease progress (**Fig. 2G**). Additionally, we find that these sensors are co-expressed with genes associated with specific signaling pathways as disease progresses (**Fig. 2C, 2D, Supplementary Fig. 2A, 2B**). For example, M_UP1 is enriched for genes related to the IL1 signaling pathway and complement cascades, whereas M_UP2 is enriched for genes of the IL6 signaling pathway and phagocytosis (**Fig. 2C, 2D, Supplementary Fig. 2B**).

MNMs also provide refinement of previously reported microglial changes during disease^18,19^. For example, down-regulation of homeostatic microglial markers previously reported in disease^18,47^, is split between two modules, M_DOWN1 and M_DOWN2, that represent distinct biological pathways (e.g. M_DOWN1 for prostaglandin synthesis and phagocytosis; M_DOWN2 for natural killer cell activation and positive cell cycle regulation; **Fig. 2C, 2G, Supplementary Fig. 2A, 2B**) and respond differently to microglial stimulation by Abeta42 and IL-4 in cell culture (**Supplementary Fig. 2C**). Lastly, we note that known common and rare disease genes occupy several different modules: M_UP1 contains *APOE, CXCR4, GRN, CSF1, PRNP, SQSTM1, TYROBP, GBA*; M_DOWN1 contains *BIN1, PVRL2, PLCG*; M_UP2 contains *TREM2, INPP5D*; M_DOWN2 contains *PICALM*; and M_UP3 contains *PSEN1* and *CD33* (**Fig. 2C**, upper panel). This annotation provides a bridge between casual disease factors and microglial stage-specific disease biology that can potentially inform our understanding of the factors that drive disease mechanisms.

These varied observations support that MNMs delineate distinct microglial transitions or states that accompany neuropathological disease progression from early neuronal dysfunction through progressive injury and neuronal cell death, building upon prior observations^18–20^ to implicate accompanying microglial functions and candidate driver genes. These modules thus provide a detailed framework for understanding phases of microglia transition related to early and later disease stages in neurodegenerative tauopathies.

### Overlap of module driver genes and transcription factors indicate substantial cross talk

Having assessed the relationship of each module to disease stage, we next moved to query the relationship of MNMs to each other. We reasoned that understanding the regulatory relationships that bridge different neuroimmune states is critical to predicting the effects of targeting these pathways for therapeutics. Using experimental gene perturbation data available through the Broad Institute’s Connectivity Map^48^ (see **Methods**), we observed that the effects of disease-related changes in MNM gene expression are not confined to the genes occupying the same MNM, but rather can effect disease-related changes of other MNMs (**Supplementary Fig 3A**). For example, perturbation of genes in the earliest up-regulated module, M_UP1, significantly upregulates genes within the subsequently up-regulated modules (M_UP2 and M_UP3) and downregulates genes within the down-regulated modules (M_DOWN1, M_DOWN2, M_DOWN3, M_DOWN4) (**Supplementary Fig 3A; Methods**). These observations that genes within the early up-regulated MNMs can drive later MNMs suggests that MNMs capture transitions from early microglial states that drive subsequent states as disease progresses.

To further delineate microglial transitions captured by MNMs, we assessed their gene promoters for shared experimentally validated transcription factor binding sites (TFBS). Nearly all MNMs showed high TFBS overlap, consistent with shared transcriptional drivers. However, two modules, M_UP1 and M_UP2 had genes with very distinct TFBS enrichments from each other (**Supplementary Fig. 3B**). This was despite substantial direct PPI connections between the modules (**Supplementary Fig. 3C**) and evidence of positive driver effects of M_UP1 genes on M_UP2 expression (**Supplementary Fig. 3A**). Therefore, while early MNMs are poised to be highly integrated at the level of regulatory drivers with later or concurrent, MNMs, M_UP1 and M_UP2 appeared to be driven by distinct candidate regulators.

### Identification of the inflammasome and anti-inflammasome related modules

To assess potential regulatory cross-talk among M_UP1 and M_UP2 genes in more detail, we re-clustered their genes to highlight any co-expression relationships that may exist between them (**Fig. 3A; Module A and Module B**). This resulted in two modules that are both up-regulated early in disease with nearly identical trajectories (**Fig. 3B, 3D**), but with strongly anti-correlated gene-module connectivity (anti-correlating kME, **Fig. 3C**), suggestive of opposing or competing pathways^26^. This is not an artifact of our transcriptional analysis, as independent CMAP gene perturbation experiments validate that gene overexpression has opposing effects on the genes clustered within these two modules (**Fig. 3F; Methods**). Other independent data confirm these relationships, in single cell RNA sequencing studies of mouse^28^ and human brain^29^ (**Fig. 3E, Supplementary Fig. 4C**), and at the level of PPI (**Supplementary Fig. 4A**). Furthermore, both modules are reproducible in independent transcriptomic datasets from mutant MAPT transgenic mice^20,35^, microglia isolated from mutant APP^37^/PS^36^ transgenic mice, and human dementia brain^20,33^, verify their robustness across mouse models of dementia and their relevance to human disease (**Supplementary Fig. 4B, 4D**). These findings support the identification of two highly conserved microglial-enriched modules that are up-regulated in disease, but that include polarized signaling pathways, which we hypothesized were poised for regulatory cross-talk.

**Figure 3:**
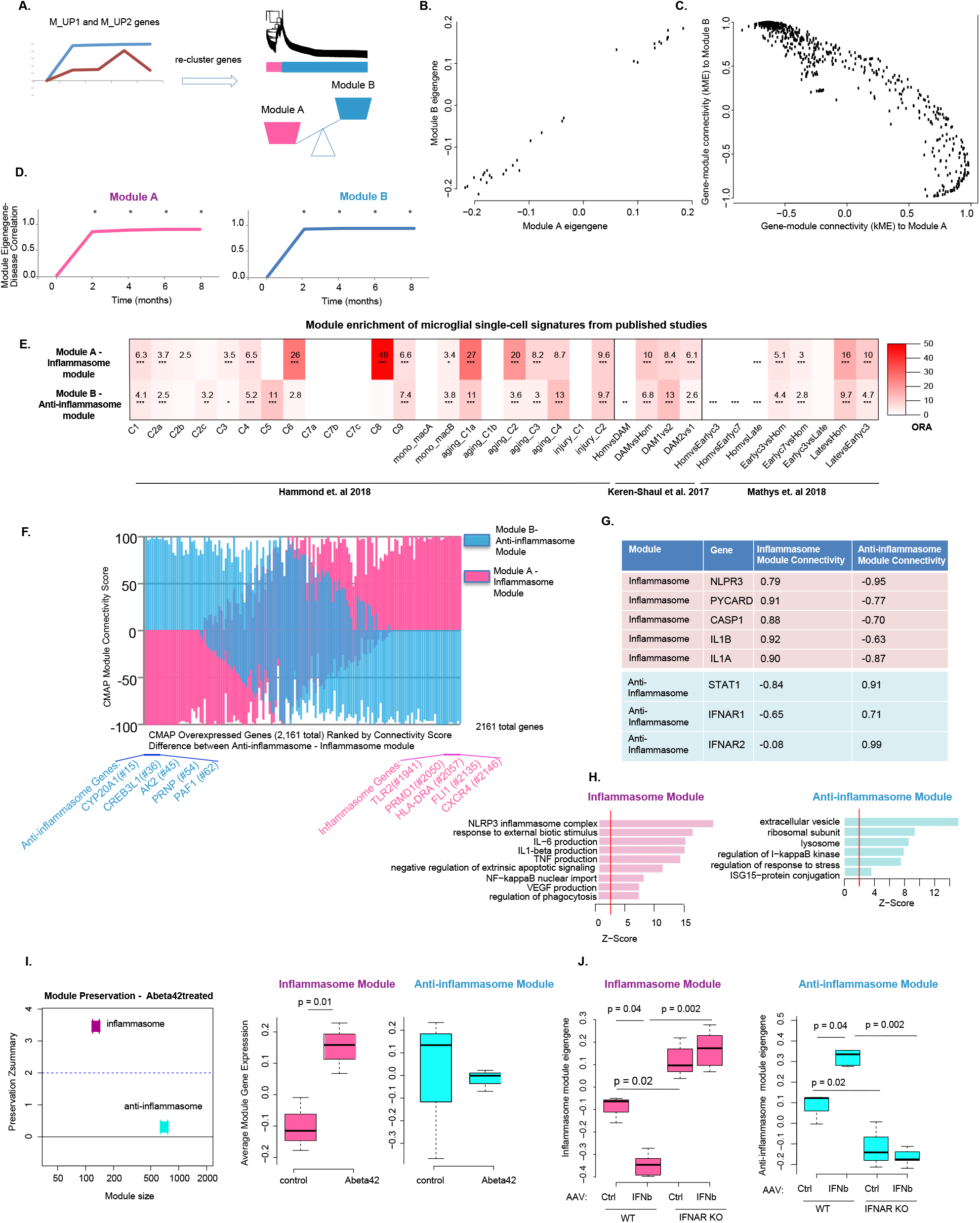
Polarized immune signaling networks are up-regulated early in disease and include signaling cross-talk among up-regulated microglia module genes. **A**, Experimental schema for identifying opposing regulatory networks among up-regulated microglia module genes. **B**, Scatterplot of module A and module B eigengenes calculated in Tg4510 purified microglia samples (n = 32). **C**, Scatterplot of gene-module connectivity scores (kME) with module A and module B calculated across Tg4510 purified microglia samples (n=32 samples, n = 899 genes). **D**, Signed Pearson’s correlation of the module eigengene (ME) calculated in the rTg4510 microglia gene expression dataset at each age (n=7 modules, n=4 mice per genotype (P301L MAPT or WT) per age, ages = 2, 4, 6 and 8 months, * two tailed p-value of Pearson’s correlation < 0.005). **E**, Module enrichment heatmap of single-cell microglial gene expression signatures from indicated published single-cell studies (Fisher’s two-tailed exact test, *FDR<0.05, **FDR<0.01, ***FDR<0.005 corrected for 2 modules and 34 total cluster signatures as defined in Hammond et al., 2018^28^, Keren-Shaul et al 2017^18^, Mathys et al. 2018^19^; Hom = homeostatic, DAM1 = type 1 disease-associated microglia (DAM), DAM2 = type 2 DAMs, as defined in^18^; from Mathys et al. 2018^19^: Hom = homeostatic (cluster 2), Earlyc3 = early response (cluster 3), Earlyc7 = early response (cluster 7), Late = late response (cluster 6)). **F**, Barplots showing CMAP connectivity scores between overexpression of a given gene (n=2161 genes) and inflammasome (pink) and ant-inflammasome (blue) modules, ordered from left to right by difference between anti-inflammasome and inflammasome module connectivity scores. Top 5 highest scoring module genes shown for each module with their ranked order among 2161 CMAP overexpressed genes. **G**, Module assignment and module connectivity scores for components of NLRP3 inflammasome complex and type 1 interferon response. **H**, Gene ontology terms significant for each module (using all module genes, permuted Z-score > 2). **I**, Module preservation and trajectory of average module gene expression of the inflammasome and anti-inflammasome modules in cultured microglia treated with oligomeric Abeta42 or vehicle control (n=3 per group, GSE55627^79^). **J**, Trajectory of inflammasome and anti-inflammasome module eigengenes in mouse microglia purified from IFNAR knockout or wild-type mice infected with IFNb expressing or control AAV (unpaired two-sample Wilcoxon rank-sum test, WT control-virus n=3, IFNAR knockout control-virus n=7, WT IFNb-virus n=5, IFNAR knockout IFNb-virus n=7, GSE98401^7^).

Module annotation and pathway analysis (**Methods**) identified the NLRP3 inflammasome and type 1-interferon response pathways as defining core components of these two modules (**Fig. 3G, 3H**), which we accordingly named the “inflammasome” and “anti-inflammasome” modules. The NLRP3 inflammasome is assembled downstream of cellular stressors and activated by the detection of various stimuli^13^, including pathological Abeta^49,50^, to promote pro-inflammatory states. Similarly, pathological Abeta rapidly and specifically stimulates the expression of the inflammasome module eigengene in microglia, both *in vivo* and *in vitro* (**Fig. 3I, Supplementary Fig. 4F**). In contrast, the prominent pathway within the anti-inflammasome module is the type-1 interferon response. Microglial isolated from mice overexpressing beta-interferon show both up-regulation of the anti-inflammasome module and down-regulation of the inflammasome module, in a manner dependent on the type 1 interferon receptor, IFNAR1 (**Fig. 3J**). Consistent with this finding, type 1 interferon is a known suppressor of the NLPR3 inflammasome^51^, and NLPR3 inflammasome activation has recently been shown to inhibit type 1-interferon signaling^52^. Furthermore, at the center of the anti-inflammasome module PPI map is MDA5 (Ifih1), a receptor of dsRNA that can activate type 1 interferon response downstream of viral detection or chromatin destabilization^44,45,53^ (**Fig. 4B**). These data provide multiple lines of evidence supporting that these two early up-regulated microglial modules represent opposing states, likely orchestrated, at least in part, by type 1 interferon signaling as a key polarizing driver.

**Figure 4:**
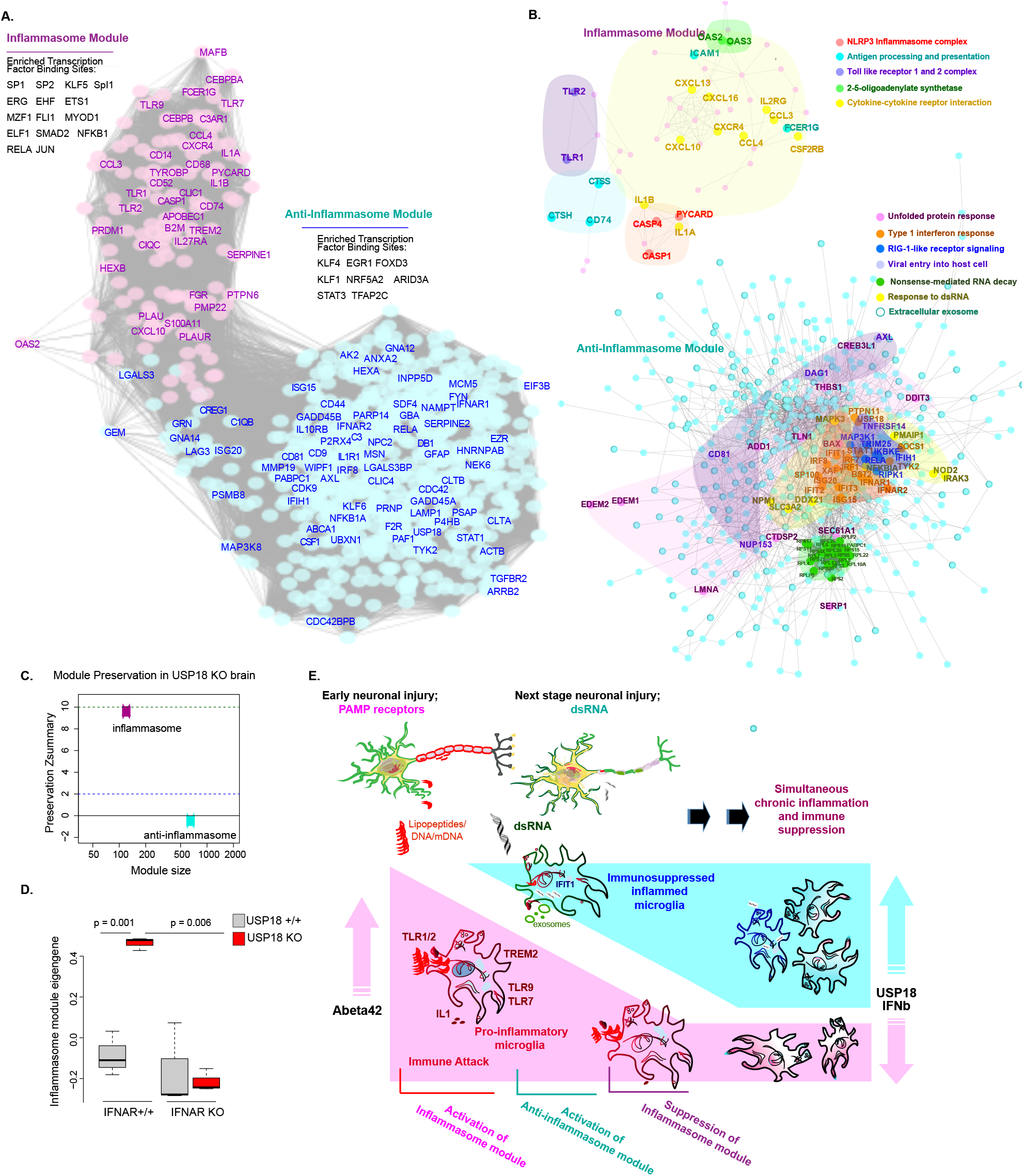
Inflammasome and anti-inflammasome modules bridge microglial sensors, mediators and checkpoints. **A**, Module gene co-expression plot of inflammasome (pink) and anti-inflammasome (blue) module genes with a distributed subset of genes labeled, and the list of transcription factors with module-wide binding site enrichment adjacent to each module (p-value relative to whole genome CpG islands < 0. 05). **B**, PPI maps with associated gene ontology pathways highlighted for the inflammasome (pink) and anti-inflammasome modules (blue). **C**, Module preservation of inflammasome and anti-inflammasome modules, and **D**, module eigengene trajectory of inflammasome module in Usp18 knockout, IFNAR1 knockout, double knockout and WT mouse brain (two-tailed unpaired T-test; n=3 per group, GSE61499^60^). **E**, Summary model of immune signaling networks represented by inflammasome and anti-inflammasome module and their interplay.

Type 1 interferon is not only a classic activator of acute anti-viral immunity, but more recently it has been demonstrated to be a critical driver of immunosuppression in the context of chronic viral infections^54,55^. Several features of the anti-inflammasome module suggest it too may represent aspects of interferon-mediated immunosuppression, including its anti-correlation with the inflammasome module, and its inclusion of genes that function as immune checkpoints (Cd274 (PDL1), Il10rb, Lag3)^54,56–58^ and inhibitors of immune activity (Usp18^59–62^, Nfkbiz, Nfkbia, Nfkbie, Tgfbr2^63,64^). Among these genes is Usp18, an established negative feedback suppressor of type 1 interferon anti-viral immune activity^59–62,65,66^ that is also highly connected with the anti-inflammasome module in microglia treated with interferon-beta (**Supplementary Fig. 4G**). Consistent with Usp18 being a critical driver of the anti-inflammasome module, we found that gene co-expression relationships in the anti-inflammasome module are completely disrupted by Usp18 knockout, without any effect on the inflammasome module (**Fig 4C**). Furthermore, the inflammasome module is highly up-regulated in the Usp18 knockout mouse in an IFNAR1 dependent fashion, suggestive of “hyperimmune” activation of the inflammasome module in the absence of Usp18 and the anti-inflammasome module (**Fig. 4D**). These data strongly support a mechanistic model, wherein interferon beta drives the anti-inflammasome and inhibits the inflammasome module through activation of immune suppressors, including Usp18, implicating interferon beta as a potential suppressor of immune activity in the chronic phase of neurodegenerative tauopathies (**Fig. 4E**), similar to what has been reported in chronic viral infections^62,65,66^.

### Viral response mechanisms link genetic risk factors across different Tau-associated dementias

Since gene expression changes on their own may represent causal, reactive or compensatory changes, we integrated genome-wide common genetic risk using MAGMA^67^ to identify whether any of the identified MNMs enrich for causal genetic factors associated with tau-related dementias. First, we identified the earliest interconnected MNM genes present in pre-symptomatic disease tissue and named them early MNM submodules, reasoning that casual disease pathways would enrich among the earliest MNM components to appear in disease (**Supplementary Figure 5A, 5B, 5E**; see **Methods**). We verified that early MNM submodules enrich for microglial signatures defined from mouse^18,19,28^ and human^29^ single cell studies (**Supplementary Fig. 5C, 5D**). We next tested all MNMs, including the early submodules, for module-wide enrichment of disease risk genes associated with FTD, AD and PSP compared to controls, based on published GWAS studies^68–70^ (see **Methods, Figure 5A**). We found that the common genetic risk associated with AD, FTD and PSP is not randomly distributed, but shows distinct patterns of enrichment: AD risk with M_UP3, FTD risk with early_UP1, FTD risk with early_DOWN1, PSP risk with early_DOWN1 (**Fig. 5A**). Each of these modules links distinct glial immune-related genes and associated pathways to disease causality, including increased exogenous antigen presentation and viral defense with AD, increased microglial immune activation and phagocytosis with FTD, and suppressed anti-viral response with both FTD and PSP (**Fig. 5B, 5C**).

**Figure 5.**
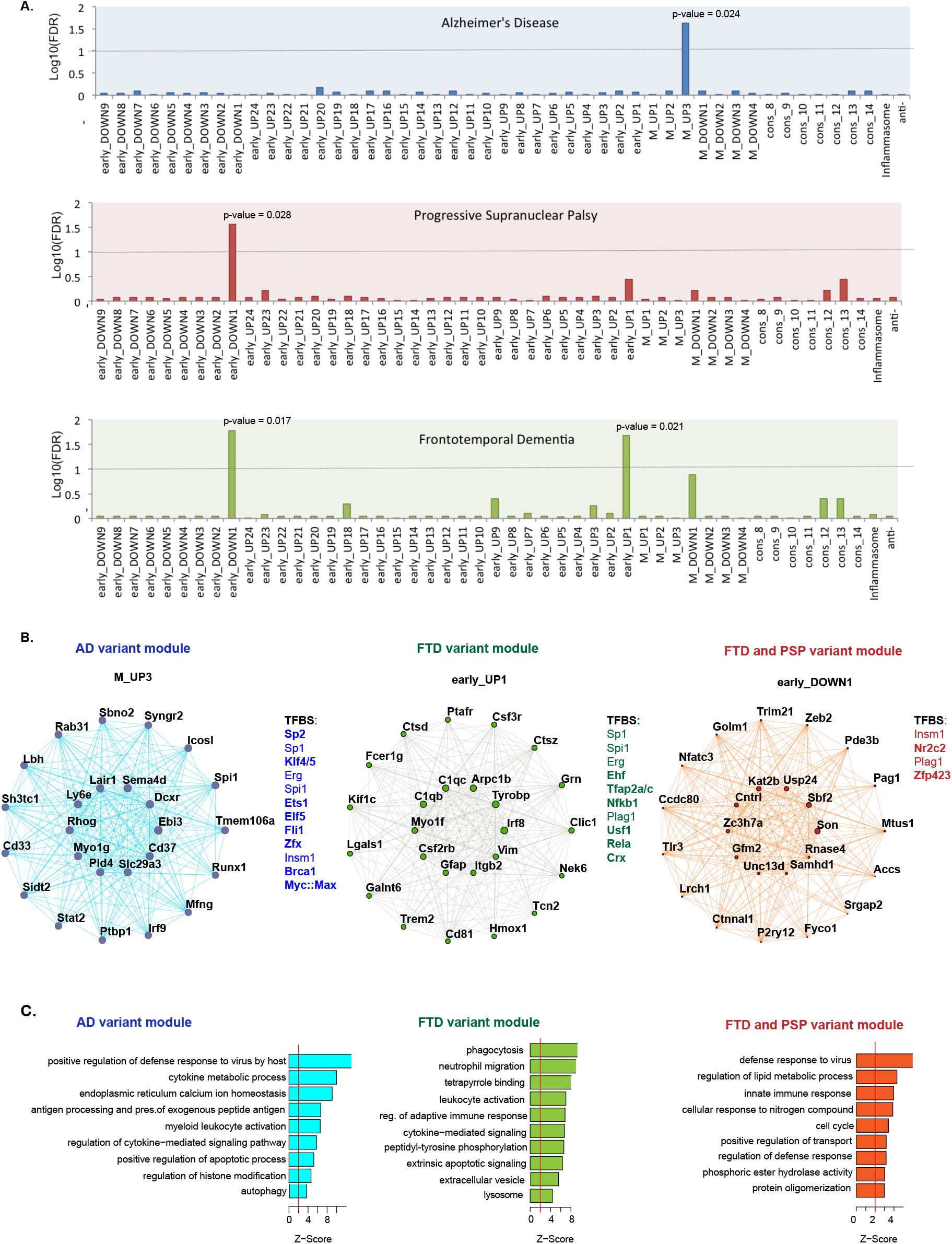
Module enrichment for GWAS variants for AD, FTD or PSP. **A**, Module enrichment for disease variants for AD^70^, FTD^97^, or PSP^69^ (FDR corrected, two-sided competitive gene-set analysis p-value from MAGMA^67^; horizontal line demarcates −log10(FDR) = 1, “anti” = anti-inflammasome). **B**, Gene coexpression network plots of top 25 genes, ranked by kME, from each disease variant-associated module, with the list of transcription factors with enriched binding sites to right of each module network plot (“TFBS”; TFs with binding site enrichment p-value <0.05 compared to whole genome CpG islands; unique TFs in bold). **C**, Gene ontology terms enrichment among corresponding module genes (permuted Z-score >2).

To independently validate these disease-module relationships, we performed confirmatory testing using a cross-disorder exome array dataset that included AD, FTD, PSP and control cases^71^. The exome array data confirmed significant associations between AD and M_UP3 (beta = 0.19 p<0.001; **Supplementary Fig. 5F**), FTD and early_UP1 (beta = 0.25, p<0.001) and FTD and early_DOWN2 (beta = 0.15 p<0.001) (**Supplementary Fig. 5F**), but not PSP, perhaps because the exome array data set is too small and therefore underpowered for PSP^71^ (**Methods**). Providing additional validation is the presence of transcription factors within these modules that are capable of inducing the disease-associated microglia gene expression patterns, including Spi1(PU.1)^72^ within the AD associated module (M_UP3) and Zeb2 within the FTD and PSP associated module (early_DOWN1) (**Fig. 5B, Supplementary Figure 5G**).

We note that viral response is a commonality among the modules enriched for common genetic variants contributing to susceptibility for these three dementias that involve tau pathology, albeit to different extents (M_UP3, and early_DOWN1) (**Fig. 5B, 5C**). This suggested a potential causal relationship between tauopathy and viral response. In the case of AD, the causal association is with late up-regulated viral response pathways, whereas for FTD and PSP the causal association is with early down-regulated antiviral response pathways (**Fig. 5B, 5C, Supplementary Figure 5E**). To test whether viral response pathways were also engaged by pathological Tau, we identified the biological pathways most correlated with Tau hyperphosphorylation in the TPR50 mouse brain (**Methods**), and indeed observed that genes involved with virus detection and anti-viral response were enriched (**Supplementary Figure 5H,I,J**), consistent with the activation of viral response pathways in concert with tau pathology in disease tissue.

## Discussion

Through an integrative systems biology approach, we have identified microglia immune networks related to specific stages of neurodegeneration modeled in mice harboring mutant Tau protein. Combining whole tissue and cell type specific data from multiple divergent mouse transgenic lines and strains, we identified seven conserved microglia modules that were also represented in post mortem tissue from patients and controls. By integrating data from brain tissue with sorted cell data, we achieved a unique perspective on neuroinflammatory signaling in neurodegeneration that we show neither can achieve on its own. Our results delineate a detailed time-course of microglial transitions across stages of progressive disease pathology that highlight specific immune receptors, biological pathways and regulatory factors at each stage. Furthermore, although each of the modules captures distinct pathways, our analysis of regulatory overlap suggest they are not entirely isolated, but rather are highly linked pathways, whose central components and core hubs transition as disease progresses through different stages. Within this robust framework, MNNs differentially implicate human disease genes with specific neuroimmune pathways that both recapitulate prior known biological relationships and identify new relationships between specific neuroimmune pathways and different disorders for further study.

Our refined analyses of microglia-associated changes across tauopathy suggest that early immune activation gives way to chronic immunosuppression, potentially driven by activation of interferon beta downstream of cytosolic dsRNA detection. In support of this is the observation that interferon beta acting through IFNAR1 is a known driver of chronic inflammatory states in cancer and chronic viral infection^54^. Furthermore, dsRNA detection can trigger interferon beta downstream of chromatin destabilization^53^. Here we find that interferon beta activates genes in the anti-inflammasome pathway capable of blocking hyperimmune activation, including Usp18^60^, and suppresses genes of the inflammasome module that participate in innate immunity. Furthermore, we find the cytosolic dsRNA receptor Ifih1 (MDA5) and associated RIG1 pathway at the core of the anti-inflammasome module PPI, and both dsRNA detection and interferon pathways to be highly correlated with pathological Tau burden (**Supplementary Fig. 5H, 5I, 5J**). These observations suggest that factors related to the accumulation of pathological Tau might trigger the interferon pathway through dsRNA detection. This is particularly salient based upon the recent observation that pathological Tau drives chromatin destabilization^73,74^, a known source of endogenous dsRNA that can activate Ifih1 (MDA5) and trigger an interferon response^44,53^. Combined, our results present a parsimonious model wherein dsRNA, released following chromatin destabilization in injured neurons in response to Tau pathology, may active chronic immune activation pathways to suppress specific immune signaling (the inflammasome module) and activate anti-inflammasome pathways to alter cellular functions including protein ubiquitination, autophagy, exosome formation, and translation^75^ (**Fig. 4E**). These observations predict that inhibition of the anti-inflammasome module, either through blockade of dsRNA, IFNAR1, or immune checkpoints within the module (PD-L1) would reduce progressive immune dysregulation triggered by pathological Tau and, at least in part, restore homeostatic microglia damage response mechanisms. These observations suggest an important causal connection between pathological Tau, viral control and the interferon response that has not previously described. Interestingly, interferon has also been implicated as a driver of microglial dysfunction in aging, suggesting interferon-driven immunosuppression in aging may also contribute to age related susceptibility to neurodegeneration^7^. Future functional and mechanistic studies will be needed to experimentally test and extend these observations, but they have potential therapeutic implications.

Genetic risk factors for AD, FTD and PSP further implicate roles for viral defense mechanisms in causal disease biology. Interestingly, the specific genes and pathways implicated differ between AD and FTD/PSP. AD genetic risk factors causally implicate antigen presentation pathways that increase in late stages in tauopathy models, whereas FTD and PSP risk factors converge upon anti-viral genes that are down-regulated in microglia very early in the mouse tauopathy models. Our findings suggest for the first time that early loss of specific anti-viral and microglial maintenance factors may be causal contributors to disease progression early in primary tauopathies.

Our findings predict that microglia may contribute to disease in a stage-specific manner by linking progressively changing disease-associated stimuli into an integrated, multi-cellular signaling network that sets the chronic course of dementia. More specifically, our observations suggest that early in tauopathy there is loss of microglial homeostasis including specific viral defense functions that promote disease progression. Further, as disease progresses and pathological Tau accumulates, it drives activation of dsRNA detection pathways, possibly through chromatin destabilization^53^, to further suppress healthy immune functions and contribute to cellular dysfunction and disease propagation. From this perspective, different stages of dementia are associated with different levels of immune activation, and as disease progresses into its clinical phase, these analyses suggest that it is likely a state of chronic immune suppression that may promote disease progression and contribute to chronic cellular dysfunction, rather than immune activation.

## Methods

### Data Set Acquisition and Filtering

Both RNAseq datasets used as input for consensus WGCNA were previously generated. The TPR50 dataset^20^ includes gene expression data from frontal cortex dissected from male mice expressing P301S MAPT or WT controls (TPR50 transgenic model^76^) in three different genetic backgrounds (C57BL6/J, F1 C57BL6/J x DAB, F1 C57BL6/J x FVB), and includes samples collected at 3 months of age (n=6 per group) and 6 months of age (n=5-6 per group). The Tg4510 microglia dataset includes gene expression data obtained from microglia purified using C11b FACS collected from mice expressing P301L MAPT and WT controls (rTg4510 transgenic model^39^), pooled to include microglia from 8-10 forebrains per sample, with n = 4 replicate samples per time points (2, 4, 6, and 8 months of age) (AMP-AD Knowledge Portal (<u>doi:10.7303/syn2580853</u>). Data were filtered for low read counts (>80% of the sample with > 10 reads with HTSeq quantification) and normalized using log2-transformation and linear regression prior to use for consensus WGCNA and module expression trajectory analysis, as previously described^20^.

Additional publicly available datasets were used throughout the study for validation or comparison. Mouse datasets consist of microarray or RNAseq transcriptomics data from a variety of transgenic mice models – Tg4510^35^, PS2APP^36^, GRN^9^, USP18^60^, IFNAR1^60^, 5xFAD^18,31^, CK-p25^19^, Zeb2^77^ and *in vitro* and *in vivo* treatments – Abeta42^78,79^, IFN-beta-expressing AAV^7^, IL4^80^, IFN-gamma^41^. Human postmortem data consist of AD temporal cortex^33^, FTD frontal cortex^20,34^, and PSP temporal cortex^33^. IRB exemption was obtained from the UCLA IRB to authorize use of de-identified human postmortem brain RNAseq data in this study.

Microarray or RNAseq datasets downloaded from the Gene Expression Omnibus (GEO) were read into R and processed as follows. Microarray data were log2-transformed and normalized by quantile normalization. Gene counts were filtered to remove low read counts (>80% of the sample with > 10 reads wth HTSeq quantification), corrected for guanine-cytosine content, gene length and library size, and log2-transformed using the CQN package in R^81^. The resulting data was used as an input to test module preservation, average gene expression and/or eigengene expression.

### mRNA Weighted Co-expression Network Analysis

In order to identify gene co-expression networks present both in purified microglia and frontal cortical brain tissue, and across multiple transgenic mouse strains and genetic backgrounds, we utilized consensus WGCNA as previously described^20^ using the WGCNA R package^26^, applied to the TPR50 dataset of forebrain RNAseq from mice aged 6 months, and the Tg4510 dataset of purified microglia (2,4,6 and 8 months), described above. The input data were generated from (1) microglia purified from P301L MAPT and WT mice from the Tg4510 model^39^ at ages 2, 4, 6 and 8 months (n=4 mice per condition) (AMP-AD Knowledge Portal (<u>doi:10.7303/syn2580853</u>), and (2) frontal cortex from P301S MAPT and WT mice from the TPR50 model with three different genetic backgrounds (C57BL6/J, F1 C57BL6/J x DAB, F1 C57BL6/J x FVB) at 6 months of age (n=5-6 per group)^20^, a period with extensive gliosis and neuronal Tau pathology but prior to frank atrophy^20^.

Biweighted mid-correlations were calculated for all pairs of genes, and then assigned similarity matrices were created using the Consensus WGCNA method as previously described^82^. In the signed network, the similarity between genes reflects the sign of the correlation of their expression profiles. The signed similarity matrix was then raised to power β to emphasize strong correlations and reduce the emphasis of weak correlations on an exponential scale. A thresholding power of 14 was chosen (as it was the smallest threshold that resulted in a scale-free R2 fit of 0.8) and the consensus network was created using the function blockwiseConsensusModules() to calculate the component-wise minimum values for topologic overlap (TOM), with parameters set as networkType = “signed”, deepSplit = 2, detectcutHeight = 0.995, consensusQuantile = 0.0, minModulesize = 100, mergeCutHeight = 0.2. Using 1 – TOM (dissTOM) as the distance measure, genes were hierarchically clustered. The resulting modules or groups of coexpressed genes were used to calculate module eigengenes (MEs; or the 1st principal component of the module). Modules were annotated using the GOElite package^83^. We performed module preservation analysis using consensus module definitions^84^. MEs were correlated with transgenic condition to find disease-associated modules. Module hubs were defined by calculating module membership (kME) values which are the Pearson correlations between each gene and each ME. Gene expression was correlated with pT231 Tau levels measured by ELISA to calculate the “gene significance” relationship with pT231 Tau, as defined by the WGCNA method^26^, using gene expression data from the TPR50 model (6 months, n=36), and this was further correlated (Pearson’s) with kME to assess the relationship between pT231 Tau and gene-module connectivity. All network plots were constructed using the Cytoscape software^85^. Module definitions from the network analysis were used to create synthetic eigengenes from which to calculate the expression trajectory of various modules in different gene expression datasets.

### Clustering of gene subsets

To apply gene co-expression methods to understand co-expression relationships among subsets of module genes in either the original consensus dataset, or in the TPR50 dataset of pre-symptomatic mice at 3 month of age, we again used the WGCNA package^26^. Biweighted mid-correlations were calculated for a subset of genes from selected consensus modules to create an adjacency matrix that was further transformed into a topological overlap matrix (with TOMType = “unsigned”). Using 1 – TOM (dissTOM) as the distance measure, genes were hierarchically clustered using the following parameters (deepSplit = 2, detectcutHeight = 0.999, minModulesize = 40, dthresh=0.1, softPower =7). The resulting modules, or groups of co-expressed genes, were used to calculate module eigengenes (MEs; or the 1st principal component of the module). The significance of intramodular connectivity was assessed for each module using a permutation test (10,000 permutations), and all modules were confirmed to have permuted p-value <0.001. “Early submodules”, described in **Figure 5** and **Supplementary Figure 5**, were derived by reclustering M_UP1 and M_UP2 genes to generate “earlyUP” modules, or M_DOWN1, M_DOWN2 and M_DOWN3 genes to generate “earlyDOWN” modules, in the 3 month of age frontal cortex TPR50 dataset previously described^20^. “Inflammasome and anti-inflammasome modules”, described in **Figure 3–4**, were derived from re-clustering M_UP1 and M_UP2 genes in the consensus WGCNA input datasets (purified microglia from the Tg4510 model and frontal cortex TPR50 dataset (6 months of age)).

### Module Preservation Analysis

We used module preservation analysis to validate co-expression in independent mouse and human datasets. Module definitions from consensus network analysis were used as reference and the analysis was used to calculate the Zsummary statistic for each module. This measure combines module density and intramodular connectivity metrics to give a composite statistic where Z > 2 suggests moderate preservation and Z > 10 suggests high preservation^84^.

### Module Gene Set Enrichment Analysis

Gene set enrichment analysis was performed using a two-sided Fisher exact test with 95% confidence intervals calculated according to the R function fisher.test(). We used p values from this two-sided approach for the one-sided test (which is equivalent to the hypergeometric p-value) as we do not a priori assume enrichment^86^. To reduce false positives, we used FDR adjusted p-values^87^ for multiple hypergeometric test comparisons. For cell-type enrichment analysis we used already published mouse brain dataset^27^. The background for over-representation analyses was chosen as total genes input into the consensus analysis (overlap of genes expressed in Tg4510 microglia and TPR50 frontal cortex RNAseq datasets).

To test module enrichment for single cell microglial gene expression signatures, we used signatures defined from published single-cell studies pertaining to microglia and/or neurodegenerative disease^18,19,28,29,77^. Specifically, for disease-associated microglia^18,19^, we set cluster signatures to be the top 100 differentially expressed genes between two microglia clusters, as defined in their corresponding publications. For microglial and macrophage clusters defined from young and aged mouse brain in^28^, we defined clusters signatures as published except duplicated genes were removed among the young cluster group (C1, C2a, C2b, C3, C4, C5, C6, C7a, C7b, C7c, C8, C9, mono_macA, mono_macB), and aged cluster group (aging_C1a, aging_C1b, aging_C2, aging_C3, aging_C4) to increase the distinctiveness of each cluster’s geneset. To define genesets from the single-cell microglial trends from injured mouse brain published in^28^, we used the genes with fold change >1.5 in control vs injured, and injured vs control mice, respectively, to define the injury_C1 and injury_C2 genesets. For human microglial gene clusters defined in^29^, we defined cluster signatures as genes with expression fold >1.8 compared to any other clusters. For Zeb2 knockout compared to control microglia, we used the published set of differentially expressed genes^77^. The background applied for over-representation analyses was set as the genes input into the consensus analysis (overlap of genes expressed in Tg4510 microglia and TPR50 frontal cortex RNAseq datasets).

### Gene set annotation

Genes in network modules were characterized using GO-Elite (version 1.2.5), using as background the set of input genes used to generated the modules being annotated^83^. GO-Elite uses a Z-score approximation of the hypergeometric distribution to assess term enrichment, and removes redundant GO or KEGG terms to give a concise output. We used 10,000 permutations and required at least 3 genes to be enriched in a given pathway at a Z score of at least 2. We report only biological process and molecular function category output.

### Protein-Protein Interaction Analysis

To assess and visualize protein-protein interactions among module genes, we used STRING (version 10.5; https://string-db.org)^88^ with the following setting (organism: *Mus musculus*; meaning of network edges: confidence; active interaction sources: experiments and databases; minimal required interaction score: medium confidence (0.400), max number of interactors to show: none). Data was exported and visualized using the Cytoscape software^85^.

### Transcription Factor Binding Site Enrichment Analysis

Transcription Factor Binding Site (TFBS) enrichment analysis using an in-house package that incorporates TFBS as previously described^89^. Briefly, we utilized TFBS position weight matrices (PWMs) from JASPAR and TRANSFAC databases^90,91^ to examine the enrichment for TFBS within each module using the Clover algorithm^92^. To compute the enrichment analysis, we utilized three different background datasets (1000 bp sequences upstream of all mouse genes, mouse CpG islands, and mouse chromosome 20 sequence). We plotted significant TFBS-module pairs (TFBS p-value < 0.05, compared to all mouse CpG islands), for TFs shared between multiple modules, as a network plot in Cytoscape, with edges connecting TFs and modules and edge weights proportional to the negative log_10_(p-value).

### Connectivity Map (CMAP) Analysis

For a given module, the top 150 module genes (ranked by kME) were used as input for the QUERY app in the Broad’s CMAP database, version CLUE (https://clue.io)^93^. This signature was used to query 7,494 gene overexpression or knockdown experiments carried out across 9 cell lines for similar (positive connectivity score) or opposite (negative connectivity score) effects on gene expression signatures, incorporating Kolmogorov-Smirnov statistics (a nonparametric, rank-based pattern-matching strategy) as described^48,93^. Mean “connectivity scores” across all cell lines was ranked by increasing order of connectivity to the input module gene expression signature to generate a rank ordered list of signed perturbagen-module connectivity scores. To identify module genes whose perturbation could reproduce the differential expression patterns of module seen in disease, we identified genes from up-regulated disease modules whose overexpression in CLUE had positivity connectivity scores with up-regulated modules or negative connectivity scores with down-regulated modules, and genes from down-regulated disease modules whose down-regulation in CLUE (via shRNA) had positive connectivity scores with signatures from up-regulated modules and negative connectivity scores with signatures from down-regulated modules, using a connectivity score cut off of |70|. Gene perturbation-module connectivity was plotted with edge length = −log10(|connectivity score|), using Cystoscope.

### MAGMA

Summary statistics for genome-wide association studies for AD^70^, PSP^69^ and FTD^68^ were used as an input for MAGMA (v1.06)^67^ for gene annotation to map SNPs onto genes (with annotate window = 20,20) and the competitive gene set analysis was performed to test module associations with GWAS variants (permutations = 100,000). All genes assigned to a given module were used as the input for each module. Consensus modules and re-clustered modules were run as separate groups in MAGMA given that they contain overlapping genes. Additional FDR correction was applied across all the competitive p-value outputs from MAGMA for all modules used in the study.

### Exome-based validation of MAGMA disease-module associations

Summary statistics from Alzheimer’s disease, Frontal Temporal Dementia and Progressive Supranuclear Palsy exome array analysis were downloaded from^71^. To incorporate protein-protein interaction, summary statistics were used as input to the network burden test, NetSig^94^. NetSig determines a gene’s network association with disease. Generalized least squares regression was used to determine if NetSig results were enriched in gene modules. Regression covariates included gene length and mean protein expression, including the log of these values. To account for linkage disequilibrium, error was correlated for genes within 5 megabase pairs.

### ELISA

Total tau and pT231 tau content were measured by commercial tau ELISA kits according to the manufacturer’s instructions (total tau – KHB0041; pT231 tau – KHB8051, Invitrogen). Briefly, standards, RIPA-soluble or sarkosyl insoluble samples were applied to the ELISA plate. After washing, a biotin-conjugated detection antibody was applied. The positive reaction was enhanced with streptavidin-HRP and colored by TMB. The absorbance at 450 nm was then measured and the concentration of tau protein was calculated from the standard curve.

## Acknowledgements

The results published here are in part based on data obtained from the AMP-AD Knowledge Portal (doi:10.7303/syn2580853). We thank Eli Lilly and Company scientists for generating the rTg4510 microglia RNAseq data and providing us access to them. For the FTD GWAS summary statistics used for MAGMA, we acknowledge the investigators of the original study (Ferrari et al, 2014, Lancet Neurol, PMID: 24943344)^68^ as well as the consortia members listed within the supplementary material. We thank Dr. Timothy Hammond and Dr. Marta Olah for use of their microglial single cell data and discussion, and Chris Hartl for helpful complementary analysis and discussion. Funding for this work was provided by Takeda Pharmaceuticals (D.H.G.), Rainwater Charitable Foundation (D.H.G.) and NIH grants to J.R (5R25 NS065723).

## Author Contribution

J. Rexach and D. Geschwind designed and supervised the experiments, analyzed and interpreted results, and wrote this manuscript. J. Rexach performed the analyses and generated figures and tables. V. Swarup contributed to preparation of raw sequencing data including quality control and normalization, and provided technical training and supervision of the cWGCNA experiments. T. Chang contributed exome-based validation of FTD, AD and PSP module associations. C. Hartl contributed to annotation of inflammasome and anti-inflammasome modules.

---

**Supplementary Figure 1.**
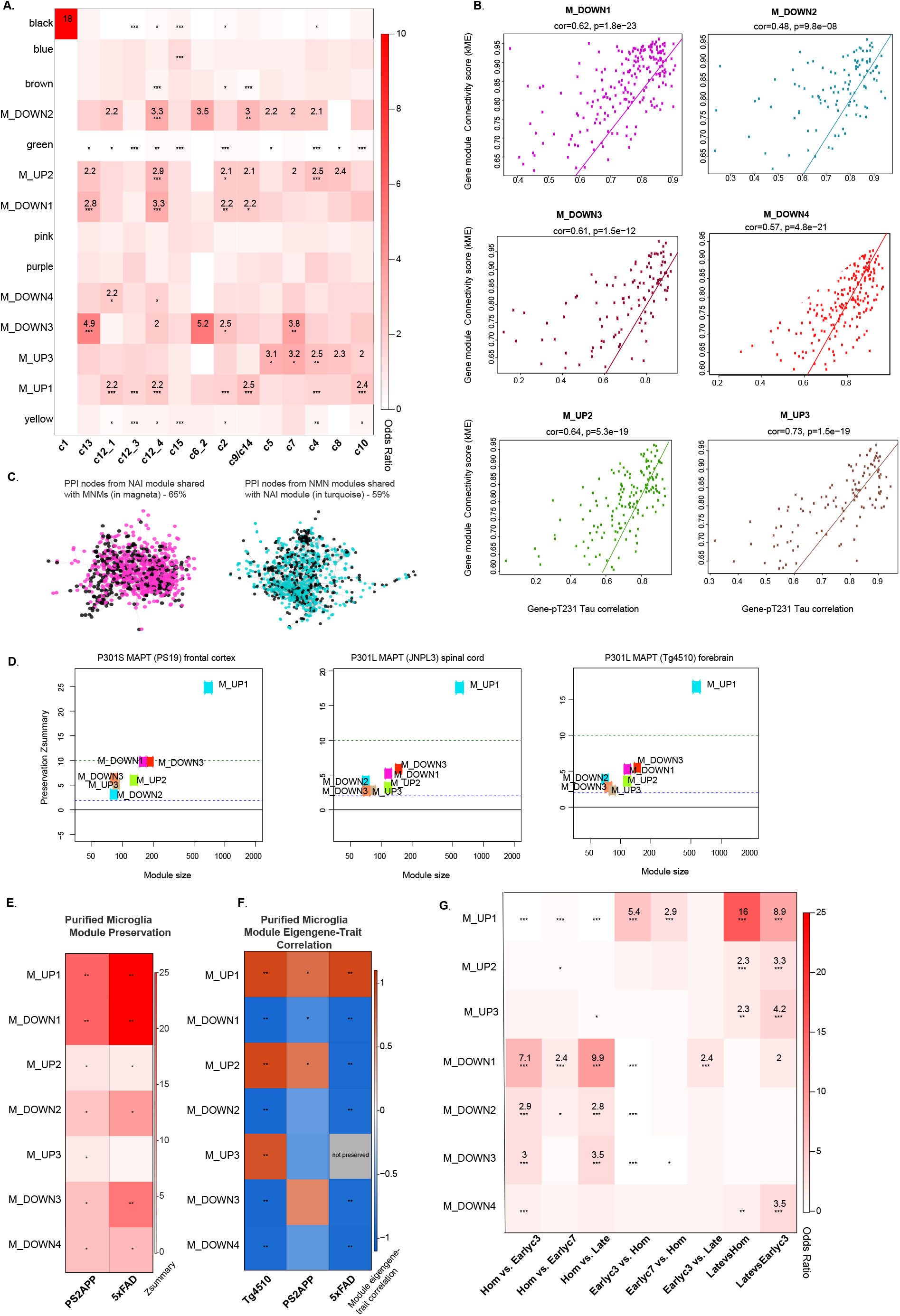
**A**, Module enrichment for human microglia single-cell cluster signatures, clusters labeled as defined in^29^ with the following abbreviations: “c1” labeled = combined c1, c3, c6_1, c11 clusters, “c13” labeled = combined c13_1 and c13_2 clusters, “c15” labeled = combined c15_1, c15_2 clusters, “c9/c14” labeled = combined c9_1, c9_2, c14_1, and C14_2 clusters (Fisher’s two-tailed exact test, *FDR<0.05, **FDR<0.001, ***FDR<0.005). **B**, Scatterplot showing Pearson’s correlation of gene-module connectivity (kME) and sample-by-sample correlation of gene expression and pT231 Tau levels (n=36; in TPR50 mouse brain; frontal cortex, 6 months of age, n=18 per group of WT or P301L MAPT; p-values obtained from two-sided test for Pearson correlation are shown). **C**, Protein-protein interaction (PPI) networks plots of the tissue-level NAI module (left) and combined MNM modules (right) showing PPI node that overlap between the two (in magenta: 65% of NAI PPI nodes overlap with MNM; and in turquoise: 59% of MNM PPI nodes overlap with NAI. **D**, Module preservation in three independent datasets of models expressing mutant MAPT: PS19 model (frontal cortex, n = 10 WT, n = 8 P301S MAPT)^20^, JNPL3 model (spinal cord, n = 12 WT, n=12 P301L MAPT, data obtained from the AMP-AD Knowledge Portal), and Tg4510 model (forebrain, n = 16 WT, n = 16 P301L MAPT; data obtained from the AMP-AD Knowledge Portal). The bottom line is at the lower cut off for preservation (Zsummary = 2) and the upper line in at the cut off for high preservation (Zsummary = 10), as defined in^84^. **E**, Module preservation heatmap in microglia purified from mouse models of Alzheimer’s pathology (5xFAD n= 5 mice per condition, GSE65067^37^; PS2APP n=5 mice per condition, GSE75431^36^). * Zsummary > 2 and <10 (low preservation) and ** Zsummary = 10 (high preservations). **F**, Module – disease trait correlation heatmap in microglia purified from mouse models of Alzheimer’s pathology (Tg4510 age = 6 months n=4 mice per genotype, 5xFAD n= 5 mice per genotype, GSE65067^37^; PS2APP age = 13 months n=5 mice per genotype, GSE75431^36^). *FDR < 0.05. ** FDR < 0.01 (n=7 modules with 3 comparisons per module) using P values from two-sided test for Pearson correlation. **G**, Module enrichment heatmap for top 100 genes differentially expressed between progressive microglia single cell states, as defined in^19^, from CK-p25 mouse model (Hom = homeostatic (cluster 2), Earlyc3 = early response (cluster 3), Earlyc7 = early response (cluster 7), Late = late response (cluster 6); n=7 modules with 9 comparisons per module, *FDR <0.05, **FDR<0.005, ***FDR<0.001).

**Supplementary Figure 2.**
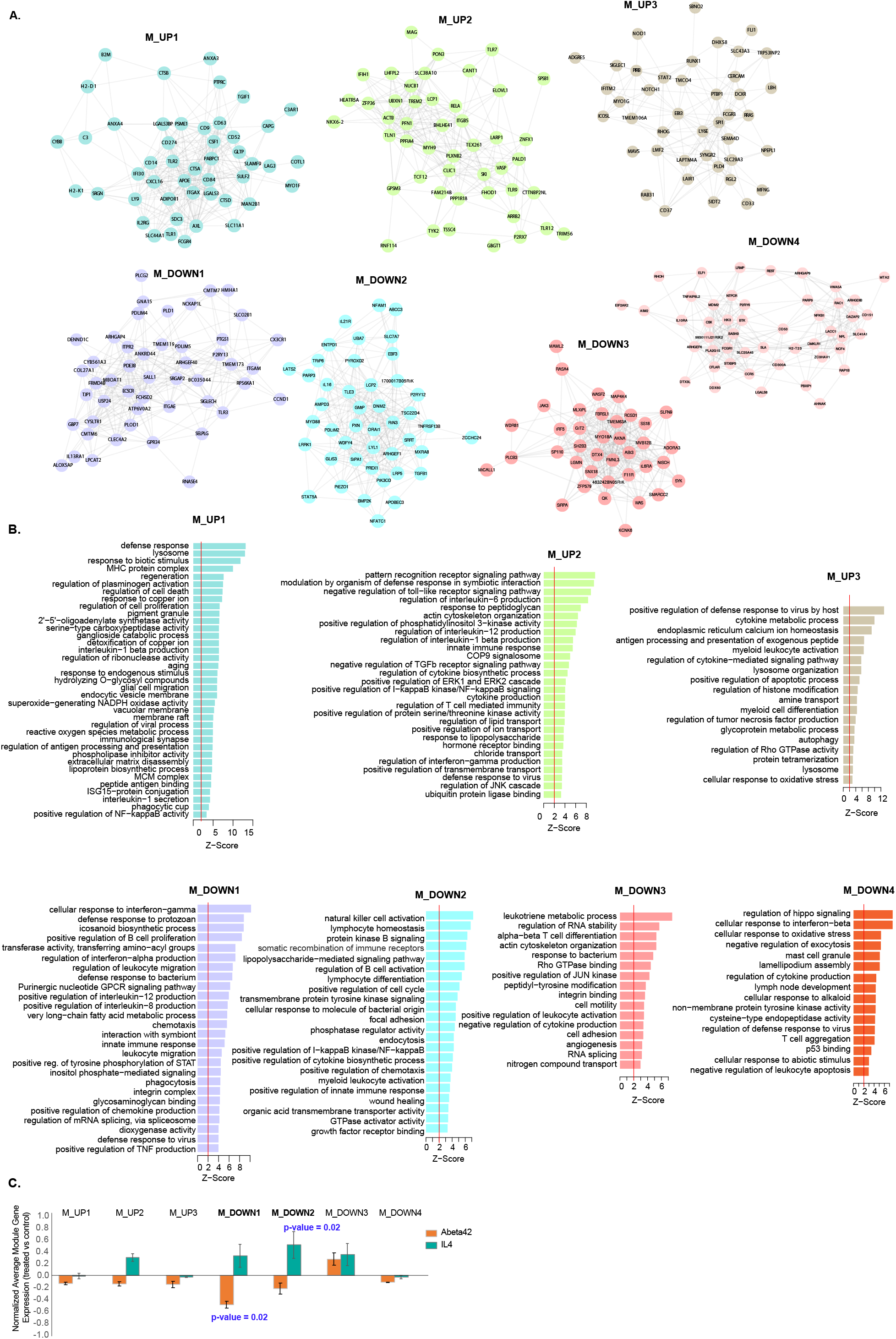
**A**, Module gene co-expression plot among top 50 module genes ranked by module eigengene connectivity (kME^26^). **B**, Extended list of gene ontology terms significant for each module (using all module genes, permuted Z-score > 2). **C**, Differential module expression in purified microglia following treatment with oligomeric Abeta42 (two-tailed unpaired T-test with FDR correction for 7 comparisons; primary mouse microglia cells, 10uM Abeta42 for 6h n=3, or vehicle for 6h n=3, GSE55627^79^) or IL-4 (two-tailed unpaired T-test with FDR correction for 7 comparisons, mouse microglia cells, IL4 at 100U/mL for 48h, n= 3, or untreated controls n=3, GSE77064^80^).

**Supplementary Figure 3.**
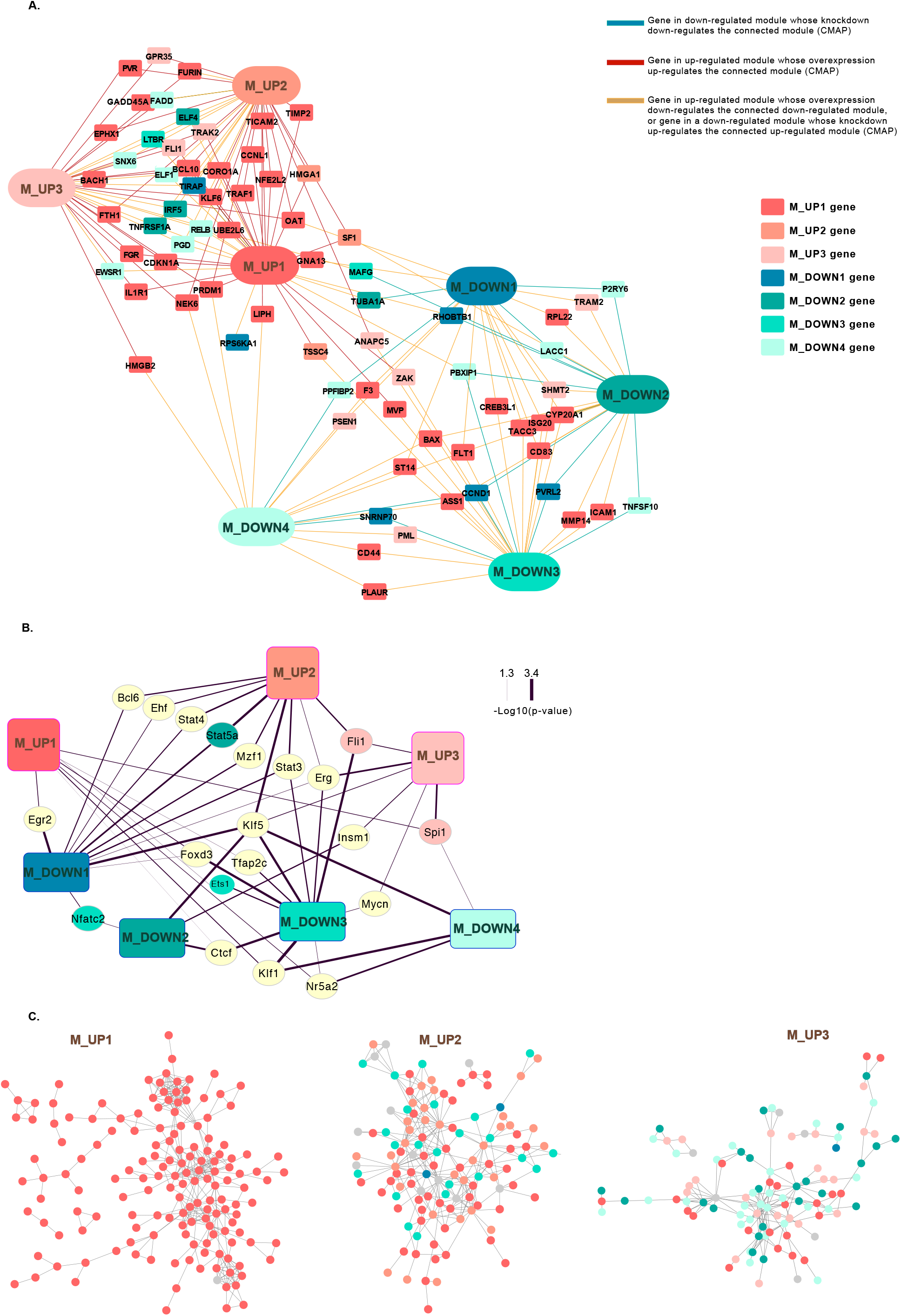
**A**, Experimental disease-associated gene perturbation-module connectivity. Connectivity between disease-associated perturbations of module genes and all modules, based on gene knockdown or overexpression experiments from CMAP, showing gene-module pairs with high connectivity (edge weighted by connectivity score-absolute connectivity score ranging 70-100- and colored by directionality of gene expression effect on module expression, as indicated). **B**, Transcription factors (TF) with binding site (BS) enrichment within more than one module (line thickness is proportion to −log10(pvalue) of TFBS enrichment within each connected module. All TFs shown have p-value < 0.05 of TFBS enrichment within module compared to genome-wide CpG islands). **C**, Protein-protein interaction maps among top 100 genes ranked by module eigengene connectivity (kME) showing later modules share protein-protein interactions with genes from other modules, including modules with TFBS overlap. Genes are colored by module, with colors as labeled in Supplementary Figure 3A and 3B.

**Supplementary Figure 4.**
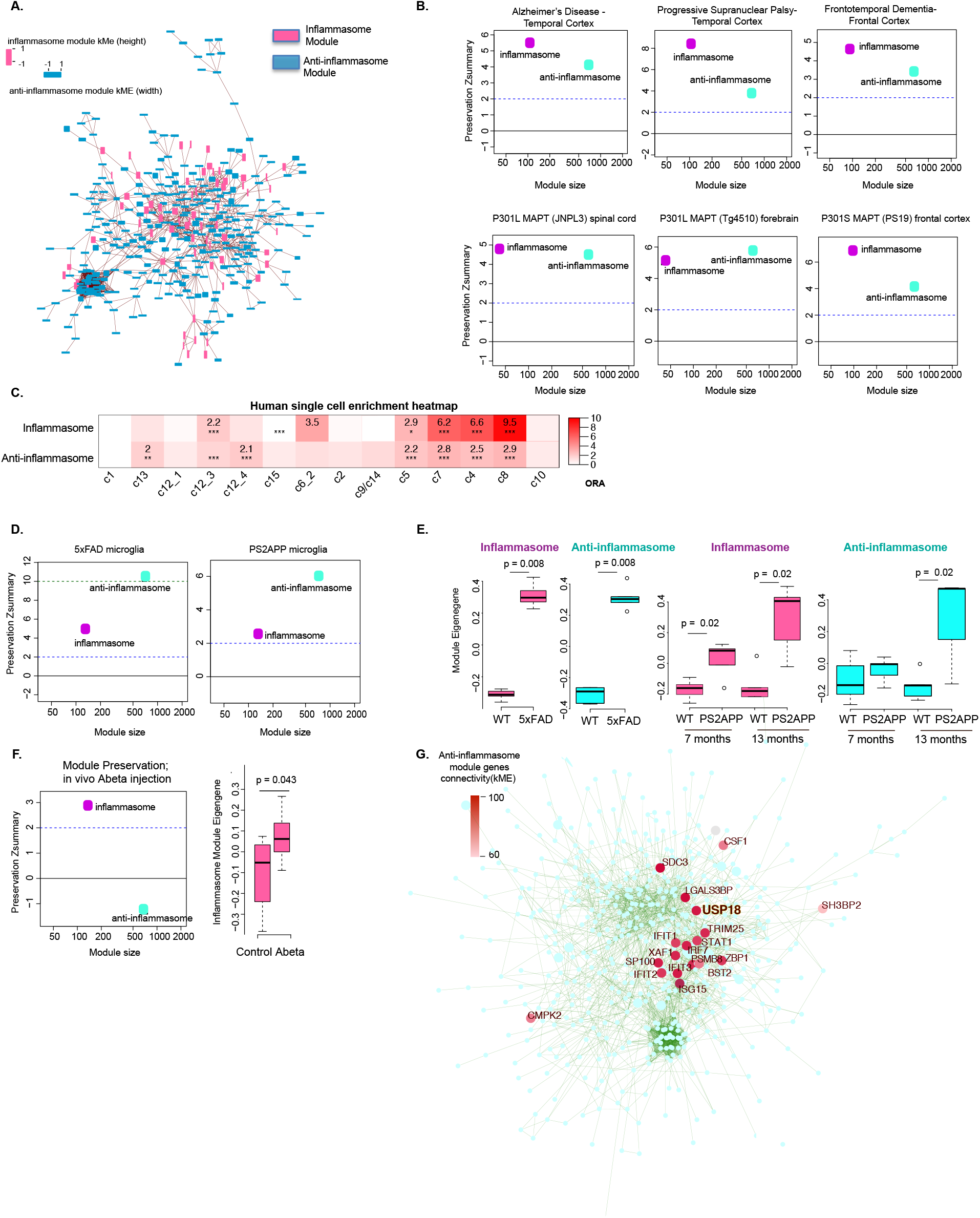
**A**, PPI network of combined inflammasome (pink) and anti-inflammasome (blue) module genes showing interconnectivity among both modules. The height and width of each node is scaled to the gene connectivity to the inflammasome (height) and anti-inflammasome (width) module eigengenes (kME^26^). **B**, Module preservation in human AD and control temporal cortex (control n=308, AD n =157)^33^, human PSP and control temporal cortex (control n=73, PSP n =83)^33^, and human FTD and control frontal cortex (control n=8, FTD n=10)^20^ and in three independent datasets of models expressing mutant MAPT: PS19 model (frontal cortex, n = 10 WT, n = 8 P301S MAPT)^20^, JNPL3 model (P301L MAPT, spinal cord, n = 12 WT, n=12 P301L MAPT, data from AMP-AD Knowledge Portal), and Tg4510 model (forebrain, n = 16 WT, n = 16 P301L MAPT, data from AMP-AD Knowledge Portal). The bottom line is at the lower cut off for preservation (Zsummary = 2) and the upper line in at the cut off for high preservation (Zsummary = 10)^84^. **C**, Module enrichment for human microglia single-cell cluster signatures, clusters labeled as defined in^29^ with the following abbreviations: “c1” labeled = combined c1, c3, c6_1, c11 clusters, “c13” labeled = combined c13_1 and c13_2 clusters, “c15” labeled = combined c15_1, c15_2 clusters, “c9/c14” labeled = combined c9_1, c9_2, c14_1, and C14_2 clusters (Fisher’s two-tailed exact test, *FDR<0.05, **FDR<0.001, ***FDR<0.005). **D**, Module preservation and **E**, trajectory of the inflammasome and anti-inflammasome module eigengenes in microglia purified from mouse models of Alzheimer’s pathology (unpaired two-sample Wilcoxon rank-sum test; 5xFAD n= 5 mice per condition, GSE65067^37^; PS2APP n=5 mice per condition, GSE75431^36^). **F**, Module preservation and module eigengene trajectory in microglia purified from 8 month old C57BL/6 mice 48 hours following injection I.C.V. with either Aβ (n=7) or vehicle (n=4) (unpaired two-sample Wilcoxon rank-sum test, GSE57181^78^). **G**, Anti-inflammasome PPI plot highlighting genes with the highest anti-inflammasome module connectivity in microglia purified from IFNb or control AAV infected mice (unpaired two-sample Wilcoxon rank-sum test, WT control-virus n=3, IFNAR knockout control-virus n=7, WT IFNb-virus n=5, IFNAR knockout IFNb-virus n=7, GSE98401^7^).

**Supplementary Figure 5.**
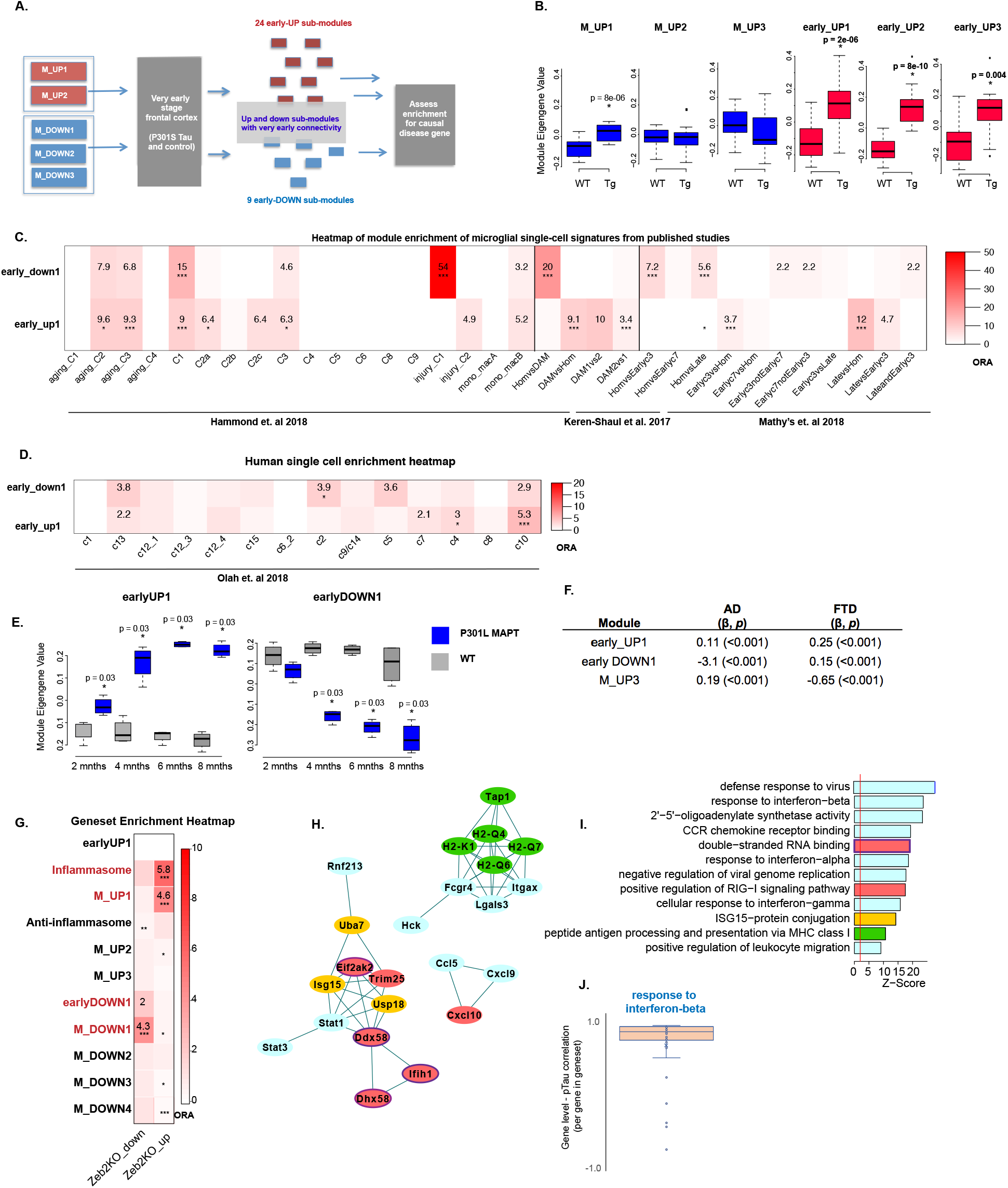
**A**, Experimental schema for identifying earliest submodules from up-regulated or down-regulated microglia module genes. **B**, Module eigengene trajectories of up-regulated module and submodules in early stage frontal cortex of TPR50 mouse model (unpaired two-sample Wilcoxon rank-sum test; age = 3 months, 3 genetic backgrounds, n=18 total P301S MAPT, n=18 WT^20^). **C**, Module enrichment heatmap of single-cell microglial gene expression signatures from indicated published singlecell studies (Fisher’s two-tailed exact test, *FDR<0.05, **FDR<0.01, ***FDR<0.005 corrected for 2 modules and 33 total cluster signatures as defined in Hammond et al., 2018^28^, Keren-Shaul et al 2017^18^, Mathys et al. 2018^19^; Hom = homeostatic, DAM1 = type 1 disease-associated microglia (DAM), DAM2 = type 2 DAMs, as defined in^18^; from Mathys et al. 2018^19^: Hom = homeostatic (cluster 2), Earlyc3 = early response (cluster 3), Earlyc7 = early response (cluster 7), Late = late response (cluster 6)). **D**, Module enrichment for human microglia single-cell cluster signatures, clusters labeled as defined in^29^ with the following exceptions: “c1” labeled = combined c1, c3, c6_1, c11 clusters, “c13” labeled = combined c13_1 and c13_2 clusters, “c15” labeled = combined c15_1, c15_2 clusters, “c9/c14” labeled = combined c9_1, c9_2, c14_1, and C14_2 clusters (Fisher’s two-tailed exact test, *FDR<0.05, **FDR<0.001, ***FDR<0.005). **E**, Module eigengene trajectory of early_UP1 and early_DOWN1 in microglia purified from Tg4510 colony (unpaired two-sample Wilcoxon rank-sum test applied at each age, P301L MAPT and WT controls, n=4 per condition per age, ages = 2, 4, 6 and 8 months). **F**, Exome-based array^71^ validation of disease-module associations identified in MAGMA (beta and p-value based on NetSig^94^ and generalized least-squares regression, see Methods). **G**, Module enrichment for genes differentially up-or down-regulated in Zeb2 knockout microglia compared to controls (Fisher’s two-tailed exact test, *FDR<0.05, **FDR<0.001, ***FDR<0.005, Zeb2 microglia data source^77^). **H**, PPI network, and **I**, enriched gene ontology terms (Z-score >2) among the top 100 genes positively correlated with Tau phosphorylation (T231) (cor > 0.9 for all top 100 genes) and **J**, box and whisker plot showing distribution, median and upper and lower quartiles of gene – pTau correlation for each gene in the GO geneset: “Interferon beta response” (GO:0035456; 74 genes), in TPR50 mouse brain (frontal cortex, 6 months of age, n=18 per group of WT or P301L MAPT).

